# Investigation of protein synthesis in Drosophila larvae using puromycin labelling

**DOI:** 10.1101/127837

**Authors:** Lisa Deliu, Abhishek Ghosh, Savraj S. Grewal

## Abstract

Translational control of gene expression is an important regulator of growth, homeostasis and aging in Drosophila. The ability to measure changes in protein synthesis in response to genetic and environmental cues is therefore important in studying these processes. Here we describe a simple and cost effective approach to assay protein synthesis in Drosophila larval cells and tissues. The method is based on the incorporation of puromycin into nascent peptide chains. Using an ex vivo approach, we label newly synthesized peptides in larvae with puromycin and then measure levels of new protein synthesis using an anti-puromycin antibody. We show that this method can detect changes in protein synthesis in specific cells and tissues in the larvae, either by immunostaining or western blotting. We find that the assay reliably detects changes in protein synthesis induced by two known stimulators of mRNA translation - the nutrient/TORC1 kinase pathway and the transcription factor dMyc. We also use the assay to describe how protein synthesis changes through larval development and in response to two environmental stressors – hypoxia and heat-shock. We propose that this puromycin-labelling assay is a simple but robust method to detect protein synthesis changes at the levels of cells, tissues or whole body in Drosophila.

## INTRODUCTION

Drosophila is an excellent genetic model system for studying animal physiology, growth, and development (Grewal, 2009; Partridge et al., 2011; Andersen et al., 2013; Padmanabha and Baker, 2014; Parsons and Foley, 2016). Over the last few decades, the versatility of Drosophila genetics has led to identification of signalling pathways and gene expression networks important for normal growth, development and aging. Moreover, the amenability of Drosophila to biochemical analyses has allowed understanding of how these networks regulate cellular biochemistry and physiology.

Regulators of protein synthesis contribute to growth, stress responses, immune responses and aging. Developing methods to measure protein synthesis in Drosophila is therefore important. Two classic methods to measure translation are polysome profiling and radioactive amino acid labelling of newly synthesized proteins. However, both have their drawbacks for analyzing protein synthesis in Drosophila – polysome profiling requires large amounts of material making it difficult to analyze specific larval cells or tissues, while radioactive amino acid labelling requires additional laboratory protocols and procedures to deal with radioactive samples. Moreover, neither approach can be used to analyze protein synthesis in situ in specific cells or tissues.

Here we present a simple, low cost assay to measure protein synthesis in Drosophila larval cells and tissues. This assay is based on a previously described puromycin labelling assay (the SUnSET assay (Schmidt et al., 2009)). Puromycin is an aminoacyl-tRNA analog that, when added to cells at low concentrations, can be incorporated into nascent peptides (Nathans, 1964; Nakano and Hara, 1979; Hansen et al., 1994). By using an anti-puromycin antibody, these newly synthesized puromycin-labelled peptides can be detected by standard immunochemical methods, and the amount of puromycin labelling hence provides a measure of nascent protein synthesis. This approach has been increasingly used to monitor protein synthesis in mammalian cells (e.g. Goodman et al., 2011; Cook et al., 2014; Dalet et al., 2017)). Here we show it can be applied to measure mRNA translational measurements in larval tissues in response to environmental and genetic manipulations.

## RESULTS AND DISCUSSION

### Measuring protein synthesis during larval development

We began by establishing conditions in which we could obtain reliable labelling of nascent peptides by puromycin. We inverted and then incubated wildtype (*w^1118^*) third instar larval carcasses in Schneider’s media containing increasing amounts of puromycin for forty minutes. We found that incorporation of puromycin increased progressively with higher concentrations of puromycin (Figure 1A). Importantly, these effects were abolished if we also co-incubated tissues with cycloheximide, indicating that the puromycin incorporation was indeed a measure of protein synthesis.

**Figure 1.**
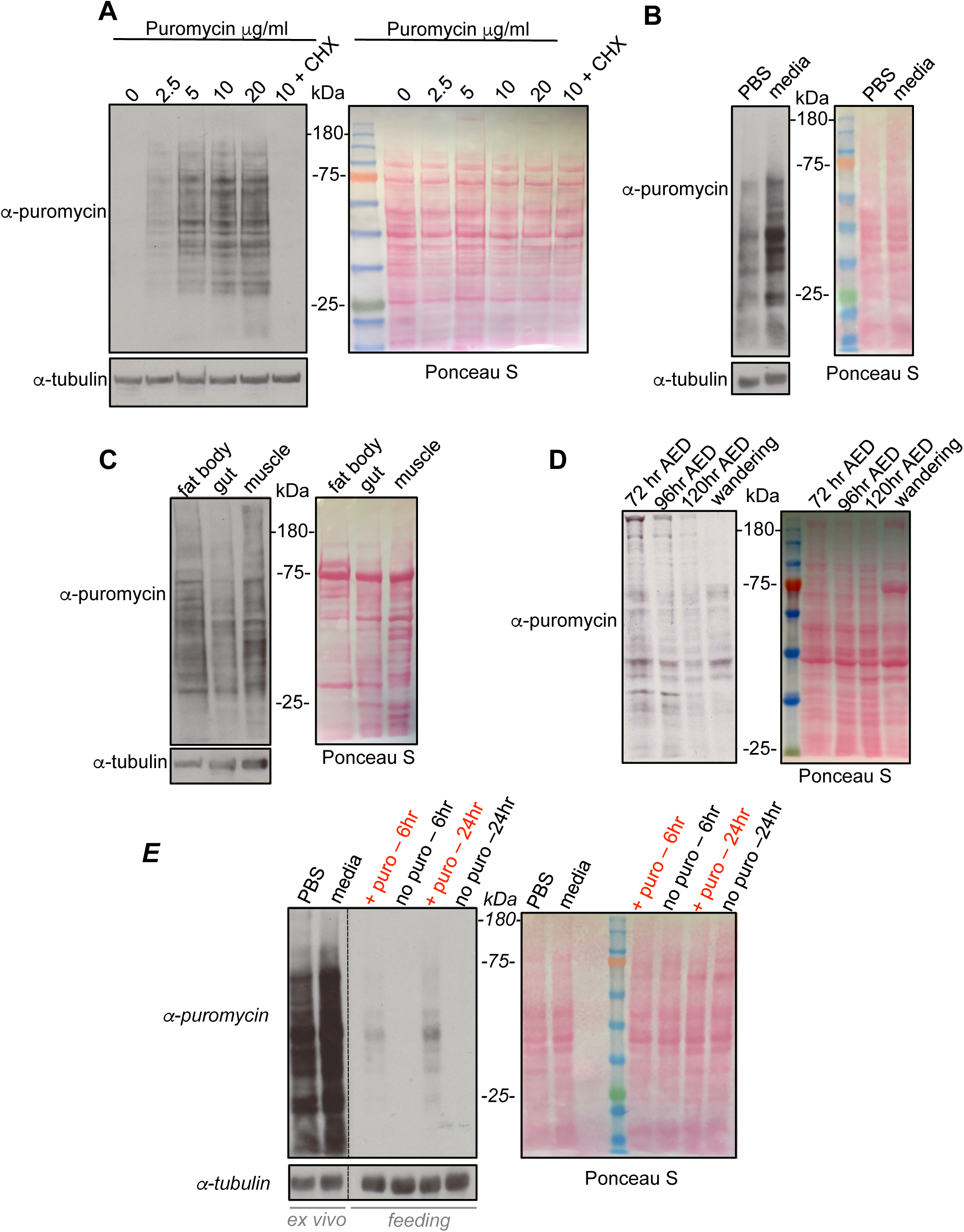
Puromycin-labelling to measure protein synthesis during larval development. (A) Larvae were inverted and whole, intact carcasses were incubated in increasing amounts of puromycin, or puromycin + cycloheximide (CHX-last lane) for 40 mins. Equal amounts of whole larval protein extracts were then analyzed by western blotting. Left, western blot with either anti-puromycin, or anti-tubulin antibodies. Right, Ponceau S staining showing total protein levels. (B) Larvae were inverted and whole, intact carcasses were incubated in either PBS + puromycin or Schneider’s media + puromycin for 40 mins. Equal amounts of whole larval protein extracts were then analyzed by western blotting. Left, western blot with either anti-puromycin, or anti-tubulin antibodies. Ponceau S staining showing total protein levels. (C) Larvae were inverted and whole, intact carcasses were incubated in Schneider’s media + puromycin for 40 mins. Larval tissues were then isolated and analyzed by western blotting. Left, western blot with either anti-puromycin, or anti-tubulin antibodies. Right, Ponceau S staining showing total protein levels. (D) Larvae at different stages in development (72hrs AED, 96hrs AED, 120hrs AED and wandering stage) were inverted and whole, intact carcasses were incubated in Schneider’s media + puromycin for 40 mins. Equal amounts of whole larval protein extracts were then analyzed by western blotting. Left, western blot with anti-puromycin. Right, Ponceau S staining showing total protein levels. (E) Comparing ex vivo vs. in vivo feeding for puromycin labelling. For the ex vivo experiments, third instar larvae were inverted and whole, intact carcasses were incubated in either PBS + puromycin or Schneider’s media + puromycin for 40 mins. For the feeding experiments, third instar larvae were transferred to either normal food (no puro) or normal food supplemented with 25 μg/ml of puromycin ( + puro) for either 6 or 24 hrs. For both the ex vivo and in vivo samples, equal amounts of whole larval protein extracts were then analyzed by western blotting. Left, western blot with either anti-puromycin, or anti-tubulin antibodies. Right, Ponceau S staining showing total protein levels. Note, the vertical dotted line in the western blots indicate where the blot was spliced to remove an empty lane and the molecular weight ladder lane (see Ponceau S staining).

We also compared the effects of carrying out the puromycin labelling in media versus PBS. We found that incorporation of puromycin did occur when larval tissues were incubated with PBS, although at a lower level than with incubation in Schneider’s media (Figure 1B). This may be because the lack of amino acids may either limit translation or my lead to loss of nutrient-dependent signaling pathways such as the TORC1 kinase pathway. It is worth noting that the levels of amino acids and glucose in Schneider’s media are approximately the same as the levels measured in larval hemolymph (Cheng et al., 2011; Pasco and Leopold, 2012). Hence, although this is an ex vivo assay, by using Schneider’s media for the labelling period, we are approximating some of the nutrient conditions in vivo.

In our assays we relied on lysis of whole larvae for our western blots. We therefore next compared puromycin incorporation in different larval tissues. We carried out the puromycin labelling as normal and then isolated specific tissues and carried out western blots. We found robust puromycin incorporation in the three tissues we tested – fat body, gut and muscle – suggesting that the assay conditions probably allow for measurement of protein synthesis in all larval tissues (Figure 1C).

We next compared protein synthesis levels at different stages of larval development. We found that protein synthesis levels were highest in larvae examined 72 hrs after egg deposition (AED) and then gradually declined throughout the remainder of larval development until wandering stage (Figure 1D).

Together these data indicate that short-term ex vivo labelling of newly synthesized peptides provides an effective way to measure translation in larval tissues and whole animals during larval development. Another potential approach is to use in vivo labelling of nascent peptides with puromycin to measure new protein synthesis. To try this, we fed larvae food mixed with puromycin and then compared the labelling of peptides by this method with the ex vivo approach described above. We initially found that feeding larvae 5mg/ml of puromycin – the amount used in our ex vivo assays – showed very little labelling, even after 24hrs of feeding (data not shown). We therefore tried a five-times higher concentration of puromycin. In this case, we did see puromycin labelling after both 6hrs and 24hrs of feeding, with the longer feeding showing higher levels of labelling (Figure 1E). However, this labelling was considerably weaker than that seen with the ex vivo method, regardless of whether the ex vivo labeling was carried out in PBS or media (Figure 1E). Hence, although feeding of puromycin provides a method for in vivo assessment of protein synthesis, it does produce much lower levels of labelling than an ex vivo method. This approach will also be subject to the caveat that altered feeding behaviour may influence the amount of puromycin-labelling.

### Effects of nutrients/TOR signaling and Myc on protein synthesis

We next examined whether the puromycin-labelling assay was sensitive to detect changes in proteins synthesis mediated by modulation of known regulators of mRNA translation. In developing larvae, the nutrient-dependent TORC1 kinase pathway is a major regulator of protein synthesis and growth (Grewal, 2009). In nutrient-rich conditions, the TORC1 kinase pathway is activated, and promotes mRNA translation and growth. However, upon nutrient deprivation, the TORC1 pathway is rapidly inhibited and protein synthesis is reduced.

We first examined the effects of six-hour nutrient deprivation on protein synthesis in third instar compared to fed controls. We carried out the puromycin labelling in Schneider’s media as above and we observed that the starved larvae showed a marked decrease on protein synthesis (Figure 2A). We reasoned that incubating the staved larvae in Schneider’s media (which contains amino acids and glucose) may potentially acutely reverse some of the physiological effects of dietary starvation. Hence we also performed the puromycin labelling by incubating fed vs. starved larval tissues in PBS plus puromycin. We saw that the overall level of protein synthesis was lower then when the assay was carried out in Schneider’s media. However, as before, we found that starvation led to a marked decrease in protein synthesis (Figure 2A).

**Figure 2.**
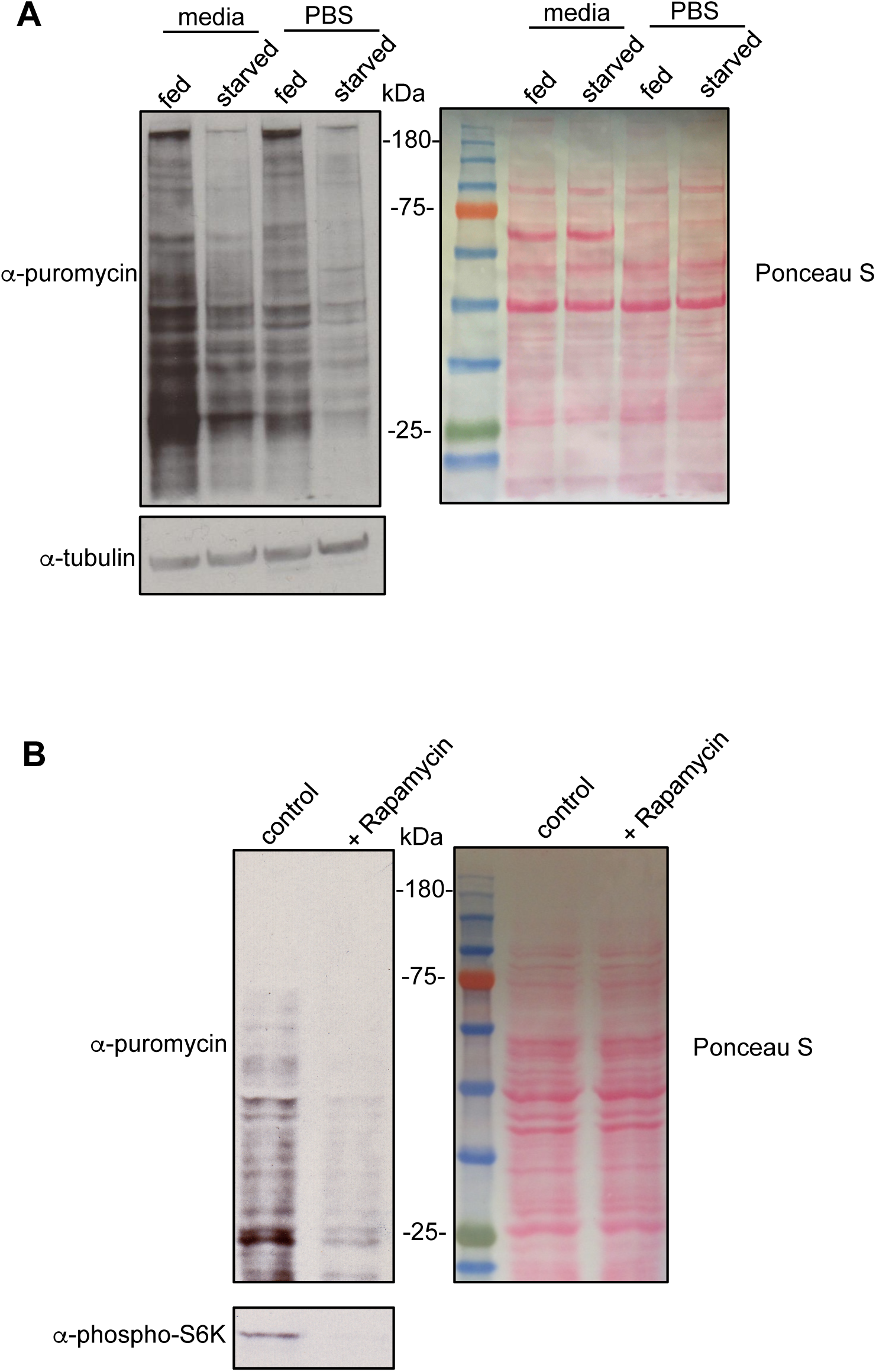
Regulation of larval protein synthesis by nutrients and TOR signaling. (A) Fed or six hour starved third instar larvae were inverted and whole, intact carcasses were incubated in either PBS + puromycin or Schneider’s media + puromycin for 40 mins. Equal amounts of whole larval protein extracts were then analyzed by western blotting. Left, western blot with either anti-puromycin, or anti-tubulin antibodies. Ponceau S staining showing total protein levels. (B) Larva were inverted and whole, intact carcasses were incubated in Schneider’s media + puromycin either with DMSO (control) or Rapamycin, 40 mins. Equal amounts of whole larval protein extracts were then analyzed by western blotting. Left, western blot with either anti-puromycin, or anti-phospho-S6K antibodies. Right, Ponceau S staining showing total protein levels.

We also looked at pharmacological inhibition of the TORC1 pathway. We carried out the puromycin labelling in third instar larvae and compared the effects of addition or absence of rapamycin – a TOR inhibitor – in the puromycin/media labelling solution. We saw that protein synthesis was markedly reduced when larval tissues were treated with rapamycin (Figure 2C).

Another regulator of protein synthesis in larvae is the transcription factor dMyc. Overexpression of dMyc increases expression of rRNA, tRNA, and ribosome biogenesis and translation factors in Drosophila (Grewal et al., 2005; Steiger et al., 2008; Marshall et al., 2012). We used the hsflp-out system to ubiquitously overexpress dMyc in third instar larvae. We found that dMyc induced a strong increase in protein synthesis compared to control animals (Figure 3A). We also tested whether the puromycin-labeling could be adapted for immunostaining to measure protein synthesis in individual cells. We used the hsflp-out system to generate GFP-marked cell clones in the larval fat body and then carried out the puromycin-labelling assay but detected puromycin incorporation by immunostaining with the anti-puromycin antibody. We found that dMyc-overexpressing fat body cells showed a marked increase in puromycin labelling compared to surrounding wildtype cells (Figure 3B). Importantly, we found that cells expressing GFP alone did not show any increase in puromycin labelling (Supplemental Fig1), suggesting that the increases in puromycin labelling did not simply reflect high levels of transgene expression.

**Figure 3.**
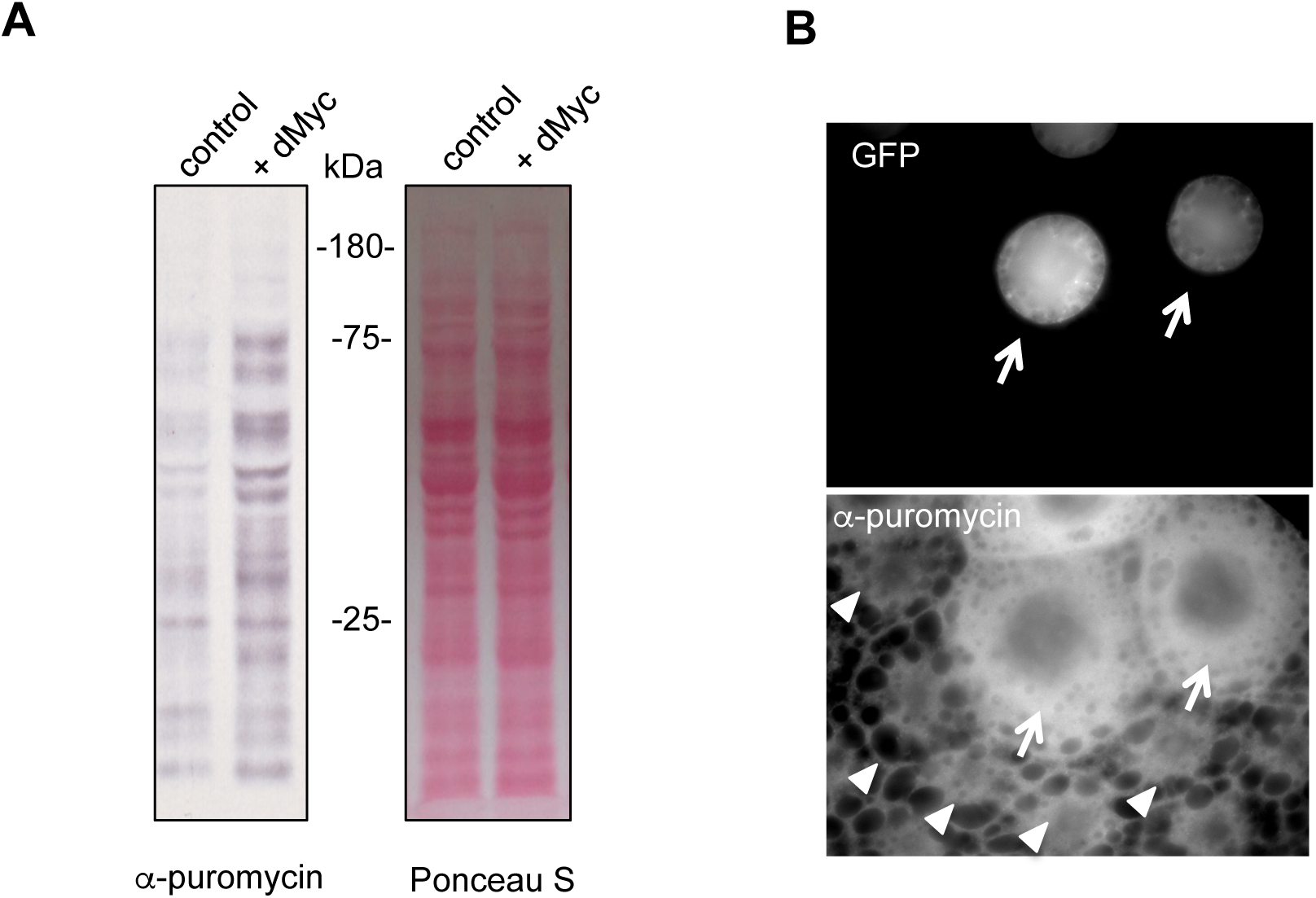
Regulation of larval protein synthesis by dMyc. (A) The hsflp-out system was used to induce ubiquitous UAS-dMyc expression in third instar larvae. Control larvae expressed UAS-GFP alone. 24hrs following transgene induction. Larva were inverted and whole, intact carcasses were incubated in Schneider’s media + puromycin for 40 mins. Equal amounts of whole larval protein extracts were then analyzed by western blotting. Left, western blot with anti-puromycin antibody. Right, Ponceau S staining showing total protein levels. Genotypes: control = *ywhsflp^122^/*+; +*/*+*; act*>*CD2*>*GAL4, UAS-GFP/*+, *dMyc* = *ywhsflp^122^/*+*; UAS-dMyc/*+*; act*>*CD2*>*GAL4, UAS-GFP/*+ (B) UAS-dMyc clones were generated in larval fat body cells using the flp-out system. Larvae were inverted and whole, intact carcasses were incubated in Schneider’s media + puromycin for 40 mins. Tissues were then immunostained with and anti-puromycin antibody. The nuclear GFP-marked cells overexpressing UAS-dMyc (arrows) show increased puromycin incorporation compared to surrounding non-GFP marked wild-type cells (arrowheads). Genotype: = *ywhsflp^122^/*+*; UAS-dMyc/*+*; act*>*CD2*>*GAL4, UAS-GFP/*+.

Together these data indicate the utility of the puromycin labelling assay to measure protein synthesis in individual cells or tissues in Drosophila larvae.

### Regulation of protein synthesis by hypoxia and heat-shock

Exposure of larvae to environmental stress has been shown to affect many conserved signaling pathways known to regulate mRNA translation in other organisms. We therefore used the puromycin labelling assay to examine how two stressors – hypoxia and heat-shock - affect protein synthesis in larvae. We first exposed third instar larvae to a 5%O_2_/95%N mixture for four hours to induce hypoxia. When we preformed puromycin labelling, we saw that the hypoxia-treated larvae showed a marked decrease in protein synthesis (Figure 4A). We next examined the effect heat-shock on protein synthesis. Third instar larvae were incubated for one hour at 37°C and then the puromycin labelling was carried out to measure their levels of protein synthesis compared to larvae maintained at 25°C. For the heat-shock conditions, we carried out the 40 min puromycin labelling at both 25 °C and 37°C. In both cases, we saw that a one hour heat-shock led to a marked increase in puromycin incorporation in a large number of peptides (Figure 4B). It is likely that many of these are members of the family of heat-shock proteins that are known to be induced by heat stress.

**Figure 4.**
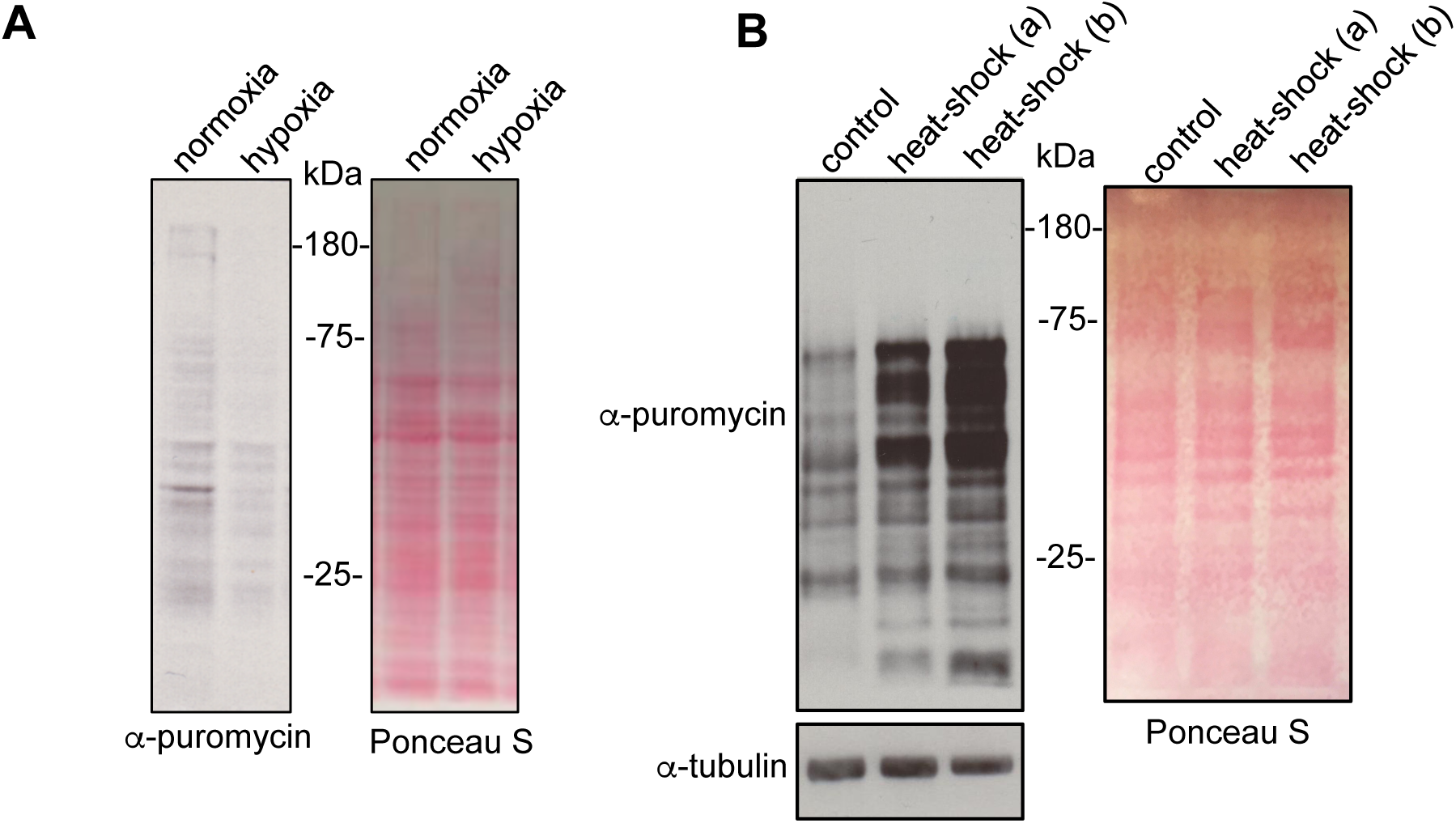
Regulation of larval protein synthesis by hypoxia and heat stress. (A) Third instar larvae were either maintained in room air (normoxia) or exposed to 5% O_2_ (hypoxia) for four hours. Larvae were then inverted and whole, intact carcasses were incubated in Schneider’s media + puromycin for 40 mins. Equal amounts of whole larval protein extracts were then analyzed by western blotting. Left, western blot with anti-puromycin antibody. Right, Ponceau S staining showing total protein levels. (B) Third instar were either maintained at 25^°^ C (control) or exposed to a one hour 37°C heat-shock. Larva were then inverted and whole, intact carcasses were incubated in Schneider’s media + puromycin for 40 mins. For the heat-shock samples the puromycin incubation was carried out either at room temperature (a) or at 37°C (b). Equal amounts of whole larval protein extracts were then analyzed by western blotting Left, western blot with anti-puromycin antibody or anti-tubulin antibody. Right, Ponceau S staining showing total protein levels.

## CONCLUSION

We describe a simple and relatively low cost ex vivo assay for robust measurement of protein synthesis in larval cells and tissues. The assay can detect both increases and decreases in protein synthesis induced by both genetic and environmental cues. Hence, the assay provides a good alternative to classic approaches to measure protein synthesis such as polysome profiling and ^35^S-methionine labelling. In addition, although Click-IT chemistry has been recently developed with modified analogs of both puromycin and methionine to measure protein synthesis in cells (Liu et al., 2012), these methods are relatively expensive compared to the puromycin labelling approach we describe here. We suggest that the ease of the assay and the ability detect translation in small amounts of tissue, will make it a useful approach to monitor how protein synthesis can be regulated in a variety of different growth, developmental and physiological conditions.

## MATERIALS AND METHODS

### Fly stocks

Flies were raised on food with the following composition: 150 g agar, 1600 g cornmeal, 770 g Torula yeast, 675 g sucrose, 2340 g D-glucose, 240 ml acid mixture (propionic acid/phosphoric acid per 34 L water. For all experiments larvae were maintained at 25°C, unless otherwise indicated. The following fly stocks were used:

*w^1118^* (used as our wild-type stock)
*ywhsflp^122^; UAS-dMyc (Grewal et al., 2005)*
*ywhsflp^122^*; +; +
*w*; +; *act*>*CD*2>*GAL*4, *UAS-GFP*

dMyc overexpression was achieved using the hsflp-out system. Early third instar larvae were heat shocked at 37°C for 2 hrs and then returned to 25°C. Puromycin assays were then carried out 24 hrs later.

### Environmental manipulations

For nutrient starvation, third instar larvae were transferred from fly food to wet filter paper and then left for six hours. For hypoxia treatments, third instar larvae were transferred to an airtight chamber perfused with a constant flow of 5% oxygen/95% nitrogen for four hours. During this period, the larvae remained in the food and were eating as normal. For heat-shock experiments, third instar larvae were transferred from 25°C to a 37°C room for one hour.

### Puromycin assay

Batches of 5-10 larvae were inverted in Schneider’s media and then the transferred to Eppendorf tubes containing media plus puromycin (Sigma). The larval samples were then left to incubate in a nutator for 40 mins at room temperature. For the experiments in Fig 1A puromycin was used at the indicated concentrations. For all remaining experiments, puromycin was used at 5μg/ml.

For drug treatments, either cycloheximide (100μg/ml) or rapamycin (20nM, Calbiochem) were added to the media/puromycin incubation solution. Following incubation, larval carcasses were snap frozen (for subsequent western blot analyses) or fixed in paraformaldehyde (for immunostaining). For experiments on specific larval tissues, at the end of the puromycin incubation period, larval carcasses were placed in ice-cold PBS and the relevant tissues were isolated and lysed for Western blot analyses.

For the puromcyin feeding experiments in Figure 1E, third instar larvae were transferred to normal food supplemented with 25 μg/ml of puromycin. Larvae were then left to feed for the indicated times (6 or 24 hrs) before being snap frozen for subsequent western blot analysis.

### Western Blotting

Larval tissues were lysed with a buffer containing 20 mM Tris-HCl (pH 8.0), 137 mM NaCl, 1 mM EDTA, 25 % glycerol, 1% NP-40 and with following inhibitors 50 mM NaF, 1 mM PMSF, 1 mM DTT, 5 mM sodium ortho vanadate (Na_3_VO_4_) and Protease Inhibitor cocktail (Roche Cat. No. 04693124001) and Phosphatase inhibitor (Roche Cat. No. 04906845001), according to the manufacturer’s instruction. Protein concentrations were measured using the Bio-Rad Dc Protein Assay kit II (5000112). For each experiment, equal amounts of protein lysates for each sample (usually 15 μg to 40μg) were resolved by SDS–PAGE and electrotransferred to a nitrocellulose membrane. Blots were then briefly stained with Ponceau S to visualize total protein and then subjected to Western blot analysis with specific antibodies. Protein bands were then visualized by chemiluminescence (enhanced ECL solution, Perkin Elmer). Primary antibodies used were anti-puromycin (3RH11] antibody (Kerafast, Catalog No. EQ0001 used at 1:1000), anti-alpha-tubulin (alpha-tubulin E7, *Drosophila* Studies Hybridoma Bank), and anti-phospho-S6K (Cell Signalling Technology).

#### Immunostaining

Following puromycin incubation, Drosophila larvae were fixed in 8% paraformaldehyde/PBS at room temperature for 45 mins. After blocking for 2hrs in 1%BSA in PBS/0.1% Triton-X 100, larval carcasses were incubated overnight in anti-puromycin antibody (1:1000). Primary antibody staining was detected using Alexa 488 (Molecular probes) goat-anti rabbit secondary antibodies. Tissues were then dissected out and mounted on coverslips using mounting media (Vectashield).

## ACKOWLEDGEMENTS

Stocks obtained from the Bloomington Drosophila Stock Center (NIH P40OD018537) were used in this study.

---

**Supplemental Figure 1.**
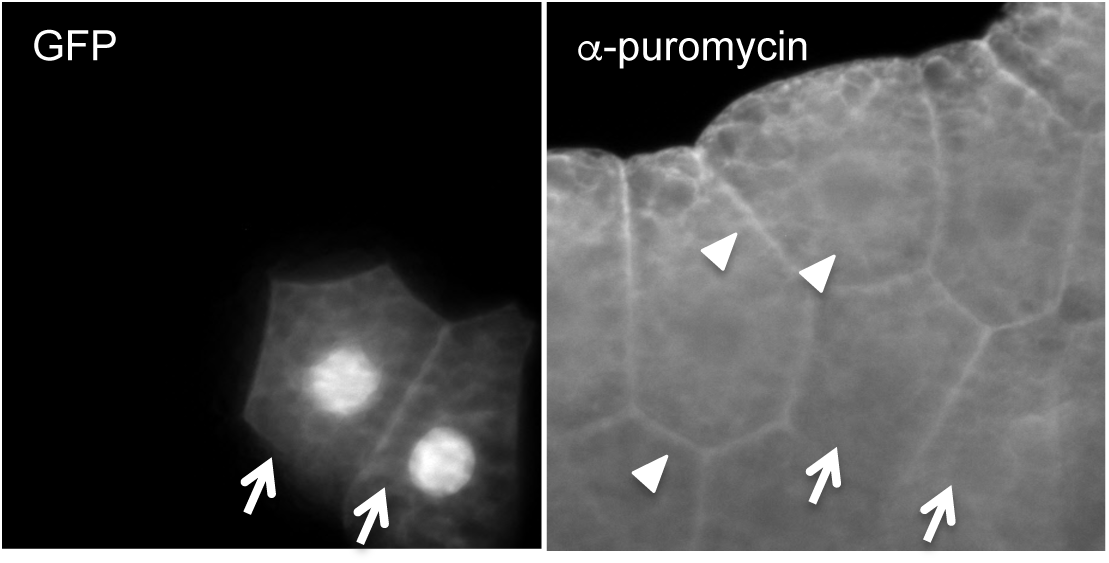
GFP expression alone does not alter puromycin incorporation. UAS-GFP clones were generated in larval fat body cells using the flp-out system. Larva were inverted and whole, intact carcasses were incubated in Schneider’s media + puromycin for 40 mins. Tissues were then immunostained with and anti-puromycin antibody. The GFP-marked cells overexpressing (arrows) show no change in puromycin incorporation compared to surrounding non-GFP marked wild-type cells (arrowheads). Genotype: *ywhsflp^122^/*+; +*/*+*; act*>*CD2*>*GAL4, UAS-GFP/*+

